# KICDB: A Causality-Oriented Multi-Omics Database for Kinase Inhibitor-Induced Cardiotoxicity

**DOI:** 10.1101/2025.08.20.671397

**Authors:** Jiamin Wei, Yin Liu, Miaoqing Wu, Guoyuan Li, Xinyao Zheng, Huafeng Fu, Jian Zhang, Jijin Lin

## Abstract

**Background:** Kinase inhibitors (KIs) are mainstays of targeted cancer therapy, but their clinical utility is frequently limited by cardiotoxicity. A systematic resource to explore the underlying causal mechanisms is urgently needed.

*Methods:* We present the KICDB (Kinase Inhibitor Cardiotoxicity Database), a comprehensive and interactive web server. KICDB is built upon a framework integrating large-scale transcriptomics meta-analysis with causal inference.

**Results:** This database centralizes the findings from a comprehensive meta-analysis of 26 kinase inhibitors (KIs) across 7 studies (n=5291) identified 8,907 significant gene expression changes in human cardiomyocytes. To establish causality, we performed a two-pronged Mendelian randomization (MR) analysis testing hundreds of downstream genes and a panel of 43 key kinase proteins against 46 cardiovascular outcomes. This large-scale analysis revealed 26 significant causal associations, implicating novel molecular mediators in KI-induced cardiotoxicity.

**Conclusions:** KICDB serves as a valuable and accessible platform for the cardio-oncology community. By integrating transcriptomic signatures with causal inference data, the database empowers researchers to formulate mechanistic hypotheses, accelerate biomarker validation, and guide the design of future cardioprotective strategies.

URL: https://zhang-lab-database.shinyapps.io/KICDB/

**Key Points:**

- We developed KICDB, a comprehensive and publicly accessible web server, to systematically investigate the causal mechanisms of KI-induced cardiotoxicity.
- KICDB integrates a large-scale meta-analysis of transcriptomic data from 26 KIs with a robust Mendelian randomization (MR) framework to move beyond correlation and infer causality.
- The analysis identified 8,907 significant gene expression changes and 26 significant causal associations between KI-associated genes and 46 cardiovascular outcomes.

## Introduction

The development of kinase inhibitors (KIs) represents a paradigm shift in oncology, offering targeted therapeutic options that have significantly improved survival rates for a multitude of malignancies ^1^. These agents, by targeting specific kinases that drive tumor proliferation and survival, offer remarkable efficacy relative to traditional chemotherapy ^2^. However, the clinical success of KIs is frequently shadowed by a significant burden of cardiovascular adverse events, a phenomenon now recognized as a major challenge in cardio-oncology ^34^. The spectrum of KI-induced cardiotoxicity is broad, encompassing hypertension, corrected QT (QTc) interval prolongation, arrhythmias, and potentially irreversible left ventricular dysfunction leading to heart failure ^5,6,7^. These complications frequently necessitate dose modifications or treatment cessation, which can compromise oncologic outcomes and increase patient morbidity and mortality ^8^.

The mechanisms underlying KI cardiotoxicity are extraordinarily complex, often stemming from both on-target inhibition of kinases vital for cardiomyocyte homeostasis and off-target effects on unintended cardiac proteins ^9,10^. A growing body of research has implicated several convergent cellular stress pathways, including the induction of mitochondrial dysfunction, activation of endoplasmic reticulum (ER) stress, apoptosis, and dysregulation of autophagic flux ^11,12^. While these studies provide crucial mechanistic clues, they are often focused on a single drug or pathway, thereby complicating efforts to construct a holistic map of KI cardiotoxicity that distinguishes class-wide effects from drug-specific ones ^13^.

A significant hurdle in translating preclinical findings to clinical practice is the methodological limitations of current research paradigms. Many mechanistic insights are derived from *in vitro* models, such as human induced pluripotent stem cell-derived cardiomyocytes (hiPSC-CMs), which, despite their utility, may not fully recapitulate the physiology of the adult human heart ^1415^. Moreover, observational studies and analyses of pharmacovigilance databases can identify statistical associations but are inherently susceptible to confounding and cannot establish causality with certainty ^16^. This critical gap between identifying molecular correlations and proving their causal role in human disease hinders the development of effective predictive biomarkers and targeted cardioprotective strategies ^17^.

To bridge this gap, we developed KICDB (**K**inase **I**nhibitor **C**ardiotoxicity **D**ata**b**ase), a comprehensive database and web server. To our knowledge, this is the first study to create a robust framework for investigating KI cardiotoxicity by integrating three distinct yet complementary analytical approaches. KICDB integrates three distinct analytical layers: (1) a systematic meta-analysis of publicly available transcriptomic data to define robust gene expression signatures; (2) a large-scale, two-sample Mendelian randomization (MR) analysis to formally test the causal relationship between these molecular signatures and clinically relevant cardiovascular diseases; and (3) functional connectivity mapping to elucidate mechanistic relationships between different KIs. By integrating these layers of evidence into an interactive, publicly accessible platform, KICDB constitutes a distinctive resource for the scientific community to move beyond correlation and explore the causal underpinnings of KI-induced cardiotoxicity.

## Materials and Methods

### Data Acquisition and Transcriptomics Meta-Analysis

To build a comprehensive transcriptomic map of kinase inhibitor (KI) cardiotoxicity, we conducted a systematic search of the NCBI Gene Expression Omnibus (GEO) database ^18^. The search was performed using the terms "kinase inhibitor" and "cardiotoxicity" to identify relevant studies. For inclusion, datasets were required to: (1) utilize human cardiomyocyte models (including primary cells and iPSC-derived lines); (2) provide raw or normalized expression data for both KI-treated and vehicle-control groups; and (3) contain clear documentation of the specific KI and treatment conditions used.

For each of the 7 included studies, raw RNA-seq count matrices were processed. Differential gene expression analysis was performed using the DESeq2 R package (v1.46.0), which models the raw counts with a negative binomial distribution to robustly identify expression changes ^19^. The primary outputs from each individual analysis—the log2 fold change (log2FC) and its corresponding standard error—were carried forward for downstream integration. For each of the 26 KIs with data from multiple studies, we performed a meta-analysis using a random-effects model, implemented in the metafor R package (v4.8-0) ^20^. This model was chosen to account for the anticipated heterogeneity in experimental systems and protocols across different studies. The final outputs were a pooled effect size (meta-log2FC) and a Benjamini-Hochberg adjusted p-value (FDR) for each gene. Genes with an FDR < 0.05 were classified as significantly dysregulated.

### Causal Inference using Two-Sample Mendelian Randomization (MR)

To advance from correlation to causation, we conducted a two-pronged, two-sample Mendelian randomization (MR) analysis using the TwoSampleMR R package (v0.6.18) ^21^. Genetic instruments (SNPs) and outcome summary statistics were sourced from the IEU OpenGWAS project database ^22^. The final set of outcomes was curated to 46 distinct cardiovascular disease phenotypes, which were grouped into 8 major categories for stratified analysis and multiple testing correction.

Our analysis tested two distinct classes of exposures: a curated panel of 43 kinase proteins was evaluated to assess direct causal effects (Kinase MR), while the top 100 most significantly dysregulated genes from each KI’s meta-analysis were selected to investigate effects mediated by downstream targets (Drug-Gene MR).

Instrument validity was rigorously enforced according to the three core MR assumptions. Relevance was ensured by selecting SNPs with a genome-wide significance (P < 5×10⁻⁸) and sufficient strength (F-statistic > 10). The Independence of instruments was achieved by pruning for linkage disequilibrium using a stringent r² threshold of < 0.01 within a 10,000 kb window. To address the Exclusion Restriction assumption, we manually screened SNPs against the GWAS ATLAS database ^23^ to remove variants with known pleiotropic effects, and further assessed this assumption using a suite of sensitivity analyses.

The primary causal estimate was derived using the random-effects inverse-variance weighted (IVW) method. To ensure the robustness of our findings, we performed multiple sensitivity analyses, including MR-Egger regression to detect directional pleiotropy, the weighted median method, and both MR-PRESSO ^24^ and MR-lap to detect and correct for outlier-driven pleiotropy. Finally, to control for the large number of tests performed, a Benjamini-Hochberg FDR correction was applied to the p-values within each of the 8 outcome categories, a strategy designed to balance statistical power with the control of false discoveries across related phenotypes.

### Functional Connectivity and Database Construction

To understand systems-level perturbations, Gene Set Enrichment Analysis (GSEA) was performed using the fgsea R package ^25^. For each KI, genes were pre-ranked by their meta-analysis log2FC to create a signature. These signatures were then queried against the MSigDB C2 (curated gene sets) collection (v2024.1.Hs) to identify significantly enriched biological pathways.

The KICDB database and web interface were constructed using the R programming language (v4.4.3) and the Shiny framework (v1.11.1) ^26^. The user interface was built with the shinydashboard package (v0.7.3) ^27^. Interactive visualizations, such as dynamic volcano plots, sortable data tables, and network graphs, were created using the plotly (v4.11.0) ^28^, DT (v0.33), and visNetwork(v2.1.2) ^29^ packages, respectively. The complete, interactive database is freely and publicly accessible at https://zhang-lab-database.shinyapps.io/KICDB/.

The R scripts used for data analysis and to build the Shiny application are available from the corresponding authors upon reasonable request.

## Results

### KICDB: A Centralized, Interactive Resource for Exploring KI Cardiotoxicity

Our study employed a multi-phase approach to construct a comprehensive resource for investigating KI cardiotoxicity, as illustrated in the study design flowchart (Figure 1a). The workflow began with the systematic collection of Data Sources, including transcriptomic data from the Gene Expression Omnibus (GEO) and genome-wide association study (GWAS) summary statistics from public repositories. This data was then processed through our Analysis Workflow, which features three primary bioinformatics modules: a transcriptomics meta-analysis to identify robust gene signatures, a functional connectivity analysis to uncover mechanistic insights, and a two-sample Mendelian randomization to infer causal associations.

**Figure 1.**
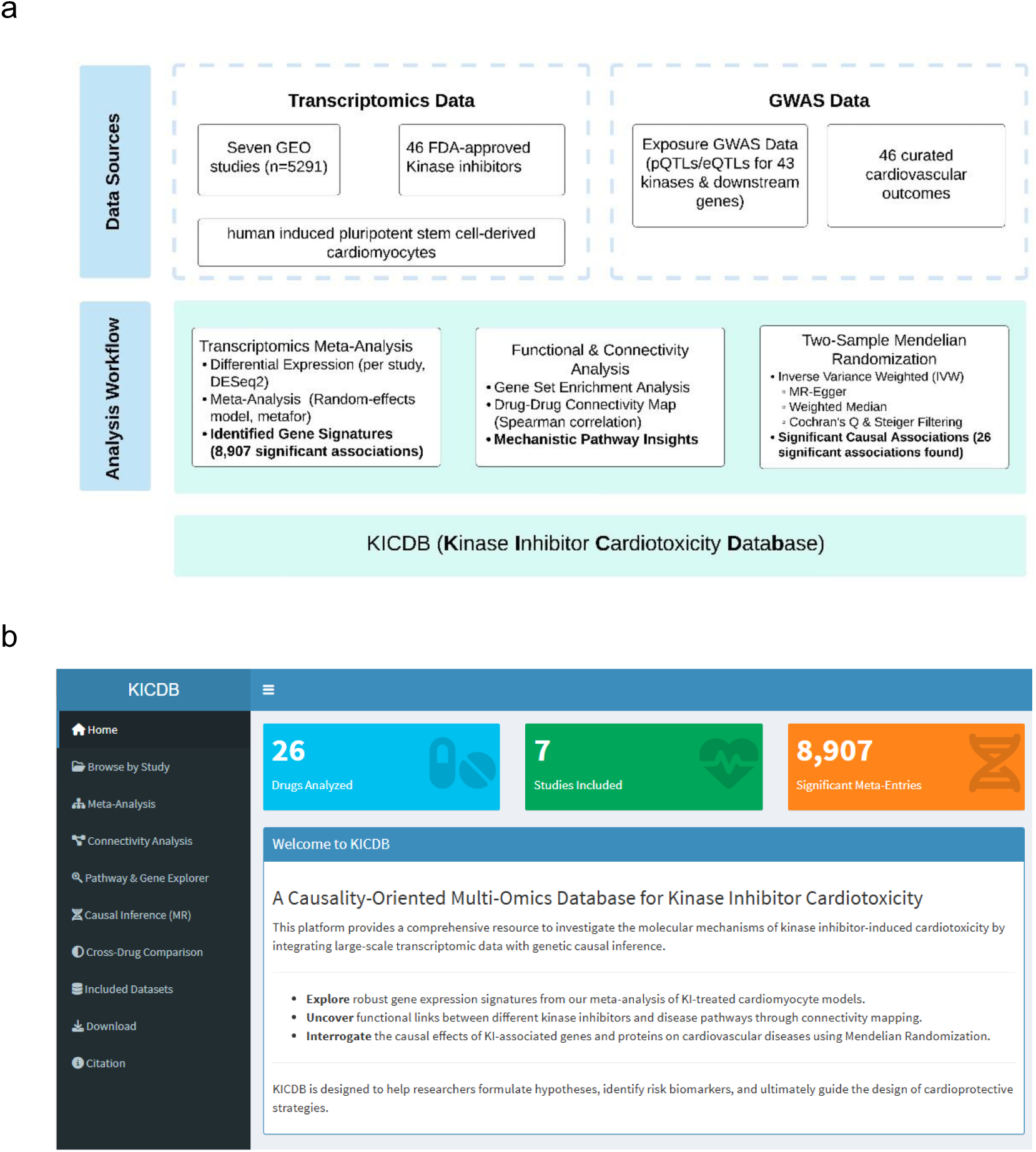
a. Overall study design, analysis workflow, and architecture of the KICDB. This flowchart illustrates the three core components of the study. All results are integrated into the final interactive KICDB Database. b. The homepage of the KICDB web application.

The culmination of this work is KICDB, an interactive web server. The homepage of the KICDB web application (Figure 1b) provides a quantitative overview of the database content, including the number of KIs, studies, and significant findings, offering users an intuitive entry point for data exploration.

### Systematic Meta-Analysis Reveals a Shared Transcriptomic Signature of Cardiotoxicity

To define the molecular landscape of KI-induced cardiotoxicity, we performed a comprehensive meta-analysis for 26 KIs across 7 independent studies. This approach allowed us to identify consistent gene expression changes while accounting for inter-study variability. As a representative example, the volcano plot of the meta-analysis for Erlotinib, Lapatinib, Sorafenib, and Sunitinib. illustrates widespread transcriptomic changes (FDR < 0.05) (Figure 2a).

**Figure 2.**
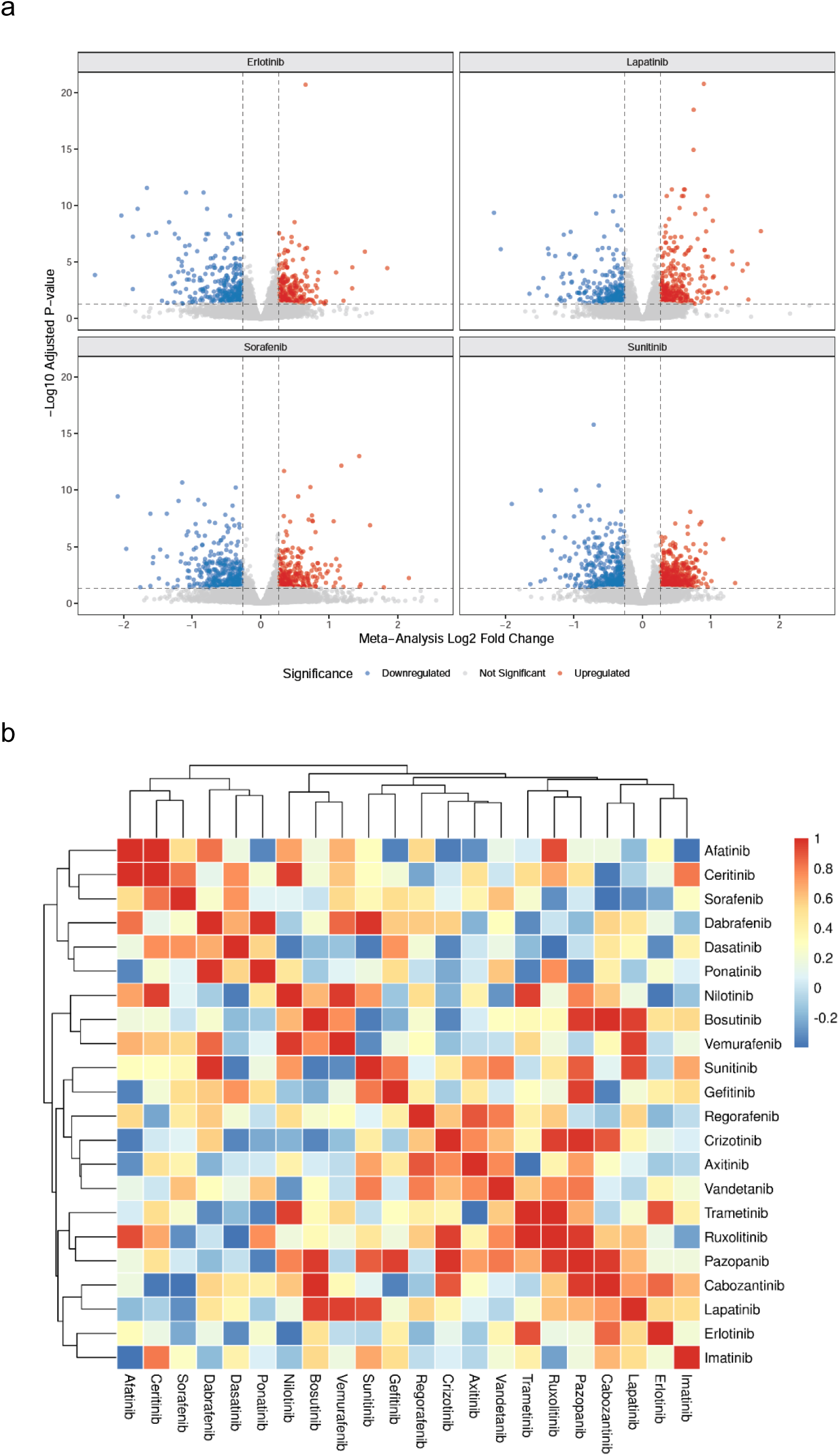
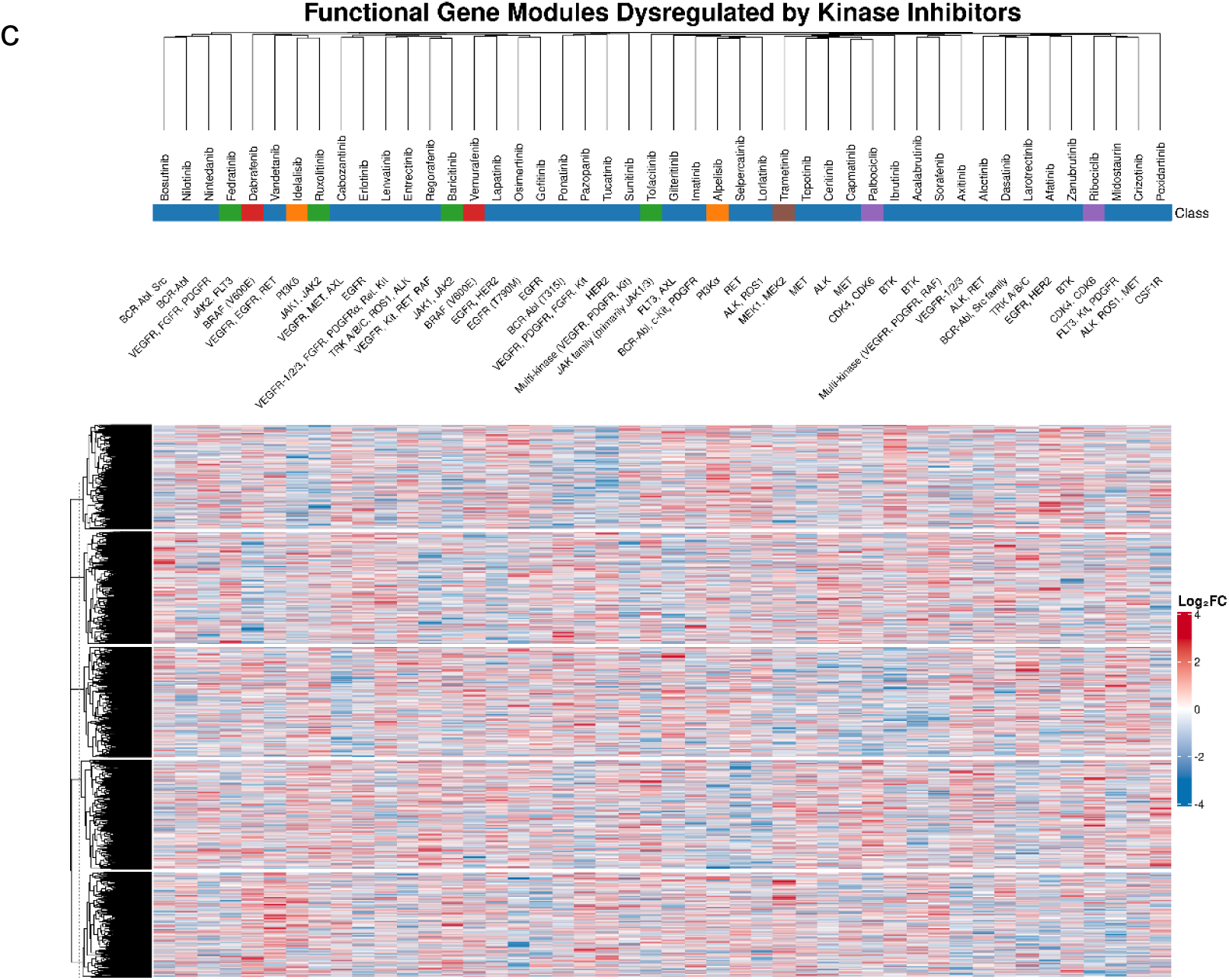
Transcriptomic meta-analysis reveals shared and drug-specific molecular signatures of kinase inhibitor (KI) cardiotoxicity. a. Volcano plot of the meta-analysis results for Erlotinib, Lapatinib, Sorafenib, and Sunitinib. Each point represents a gene, with red and blue points indicating significantly up- and down-regulated genes (FDR < 0.05), respectively. b. Heatmap of Spearman correlation coefficients calculated from the log2FC profiles of 26 KIs from the meta-analysis. Hierarchical clustering groups drugs with similar transcriptomic effects. c. Hierarchically clustered heatmap of log2FC values for significant genes (rows) across 26 KIs (columns). Top annotations show drug class and targets; row annotations list the top Gene Ontology (GO) term for each of the five gene modules. Red/blue cells indicate gene up/down-regulation.

To assess the global similarity of these effects across all KIs, we performed hierarchical clustering on a Spearman correlation matrix of their expression profiles. The resulting heatmap revealed distinct clusters of drugs with similar transcriptional programs, such as the close relationship observed between Sunitinib and Sorafenib (Figure 2b).

To further dissect the gene expression patterns driving these relationships, we visualized the significantly dysregulated genes in a comprehensive heatmap (Figure 2c). Hierarchical clustering of both drugs (columns) and genes (rows) identifies distinct patterns of expression changes. The top annotation bar categorizes each drug by its primary mechanism and lists its main protein targets. This analysis partitioned the genes into five functional modules, with the top enriched Gene Ontology term for each module listed on the left, revealing how different groups of KIs converge on distinct biological pathways such as cellular organization and metabolic processes.

### Mendelian Randomization Establishes Causal Links Between KI-Associated Genes and Heart Disease

Using a two-sample Mendelian Randomization (MR) framework, we established causal links (P < 0.05) between several kinase-associated genes and cardiovascular diseases (CVDs). Our primary findings, including odds ratios (ORs) and 95% confidence intervals (CIs), are summarized in Table 1. Our investigation revealed that genetic predisposition to certain genes is causally linked to a significantly increased risk of specific cardiac conditions. A particularly strong association was discovered for RNF13, where genetic variants predicting higher expression were linked to a 37% increased risk of primary cardiomyopathies (OR=1.37) and a 12% increased risk of coronary atherosclerosis (OR=1.12). In parallel, genetically predicted higher levels of soluble TIE1 were causally associated with a heightened risk of myocardial infarction (OR=1.03) and a notable 19% increased risk of general cardiovascular disease (OR=1.19), highlighting its potential role as a risk factor across a spectrum of heart-related pathologies.

**Table 1.**
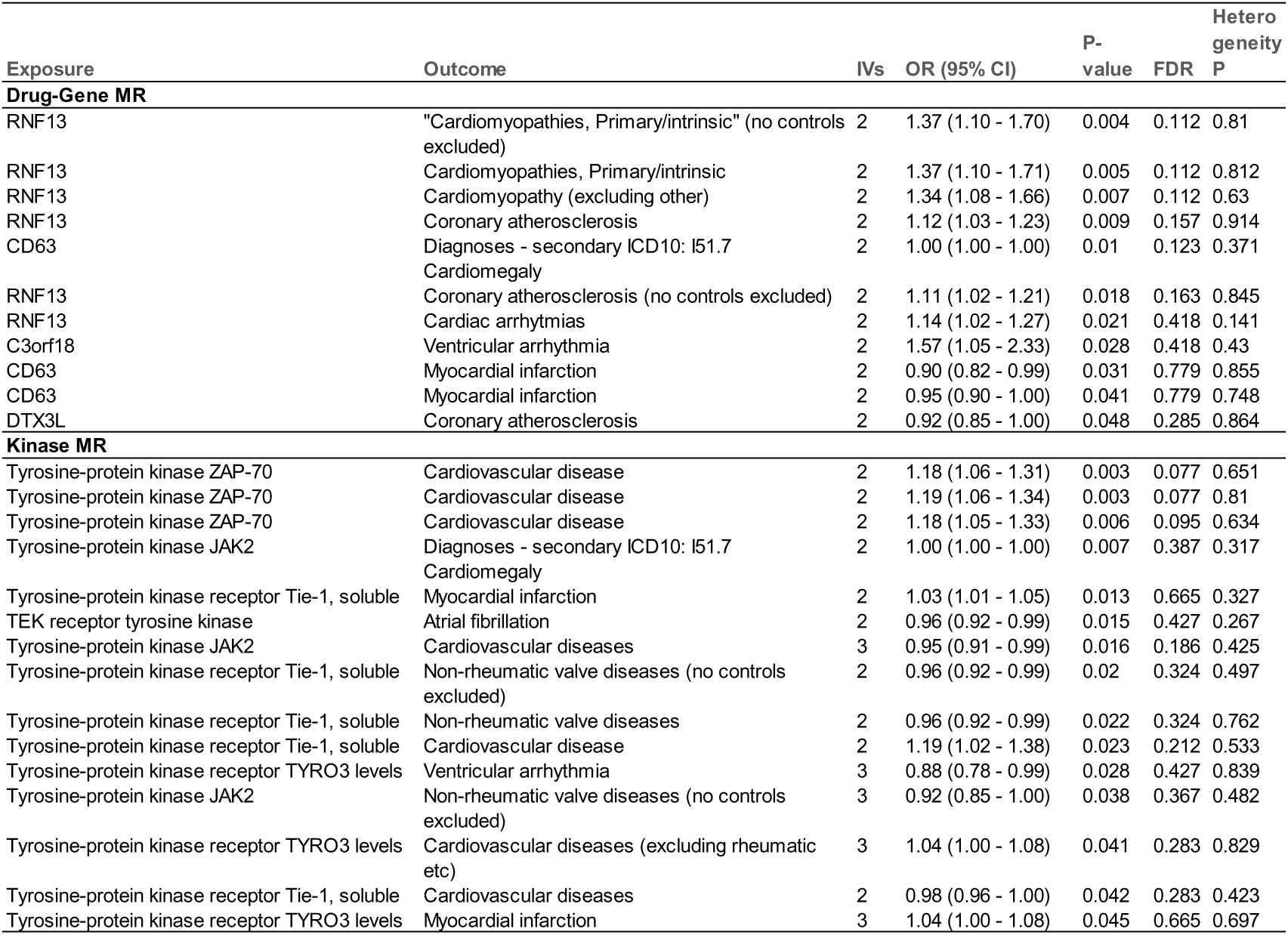
Causal Estimates from Mendelian Randomization of Kinase-Associated Gene Exposures on Cardiovascular Disease Outcomes. The table presents the results from a two-sample Mendelian randomization analysis linking genetically predicted levels of kinase-associated exposures to various cardiovascular outcomes. Only statistically significant or notable findings are shown. Abbreviations: IVs, number of instrumental variables; OR, odds ratio; CI, confidence interval; FDR, p-value adjusted for false discovery rate. The Heterogeneity P-value tests for heterogeneity among the instrumental variables, with P > 0.05 indicating no significant heterogeneity.

Conversely, our analysis also provided strong genetic evidence for several genes that exhibit a protective effect against CVDs. We found that genetic predisposition to TEK receptor tyrosine kinase was causally associated with a 4% reduced risk of atrial fibrillation (OR=0.96), suggesting a role in maintaining normal cardiac rhythm. Furthermore, JAK2 was linked to a 5% reduced risk of general cardiovascular diseases (OR=0.95), while TYRO3 demonstrated a substantial protective effect, associated with a 12% reduced risk of ventricular arrhythmia (OR=0.88). These findings pinpoint specific biological pathways that may confer cardiovascular resilience.

The validity and reliability of these causal estimates are substantially strengthened by our robustness checks. All reported significant associations successfully passed tests for heterogeneity (P > 0.05), indicating that the instrumental variables used in the analysis were not subject to significant pleiotropy, a potential source of confounding in MR studies.

### Functional Connectivity Mapping Uncovers Mechanistic Similarities Among KIs

To systematically map the functional landscape of kinase inhibitors (KIs), we constructed an undirected drug-drug functional association network (Figure 3a). This network is built from statistically significant associations (padj < 0.05) derived from connectivity mapping analysis. The final visualization employs the Fruchterman-Reingold layout algorithm to spatially organize the nodes (drugs). In the network, the edges connecting two drugs represent a significant functional relationship. The edge color is mapped to the Normalized Enrichment Score (NES), where red (firebrick2) indicates a positive correlation (synergistic or similar mechanisms) and blue (steelblue) denotes a negative correlation (antagonistic mechanisms). The edge width is directly proportional to the statistical significance (-log10(P.adj)), making stronger associations visually prominent. For instance, the thick red edge between Nilotinib and Lapatinib highlights a highly significant positive functional association, suggesting they act through convergent downstream pathways.

**Figure 3.**
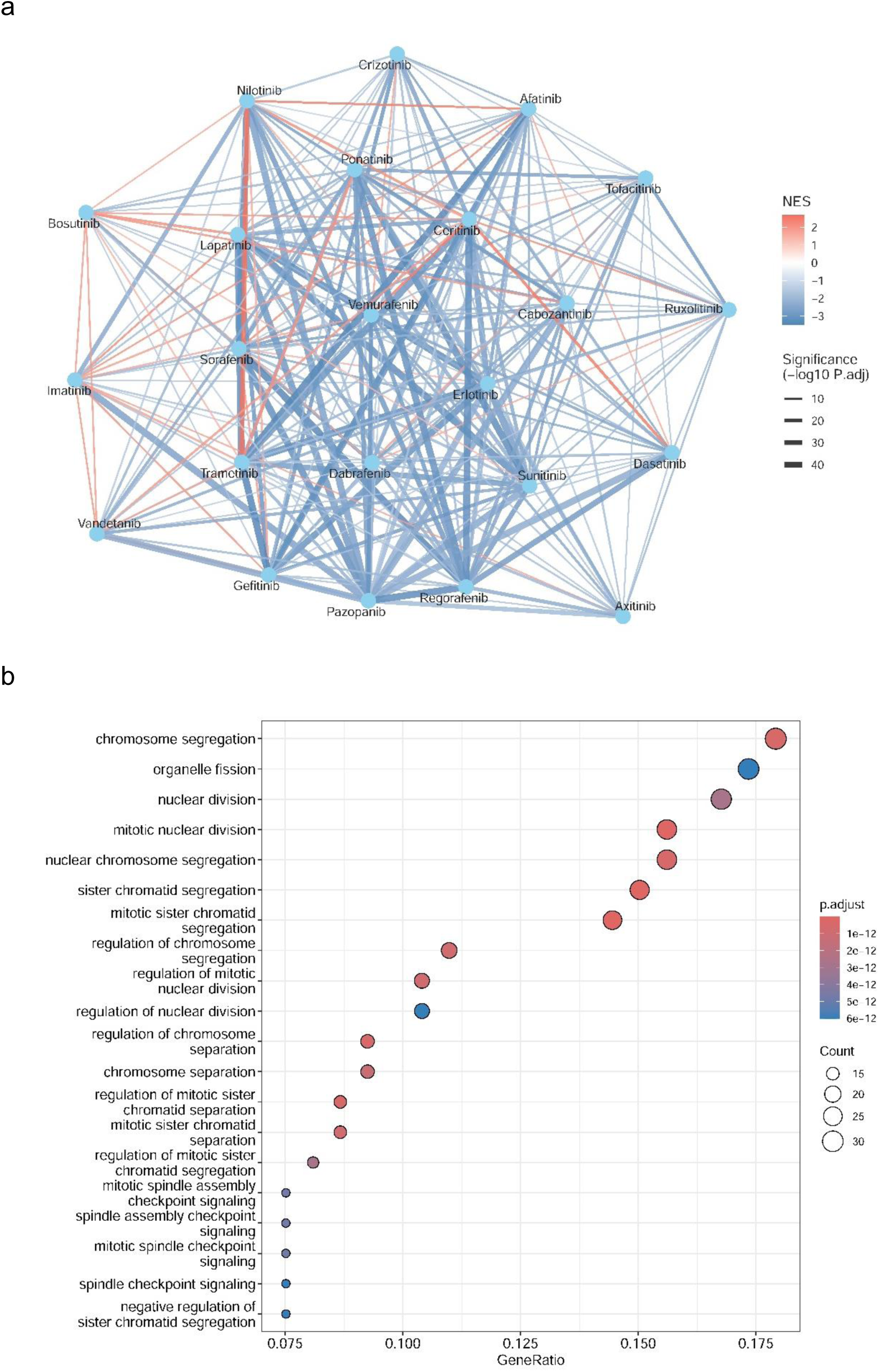
a. Drug-Drug Functional Association Network. The network shows significant functional associations between kinase inhibitors (KIs). Nodes represent KIs. Edge color corresponds to the Normalized Enrichment Score (NES; red for positive, blue for negative correlation), and edge width represents statistical significance (−log10(P.adj)); b. Enriched GO Pathways for Common Genes. Results of a Gene Ontology (GO) enrichment analysis for Biological Processes, performed on genes commonly dysregulated across at least five KI datasets. The top 20 enriched pathways are shown. Dot color indicates significance (adjusted p-value), size corresponds to the gene count, and the x-axis represents the GeneRatio.

To identify the core biological processes commonly modulated by these KIs, we performed a Gene Ontology (GO) enrichment analysis for Biological Process (BP) terms (Figure 2b). The input for this analysis was a set of "common genes," defined as those found to be significantly dysregulated in the datasets of at least five different KIs. The resulting dot plot displays the top 20 most significantly enriched GO terms, ranked by their Benjamini-Hochberg adjusted p-value. The plot clearly shows that these common genes are overwhelmingly enriched in pathways central to cell division, such as "chromosome segregation," "organelle fission," "nuclear division," and "mitotic sister chromatid segregation." These findings strongly suggest that a primary, shared mechanism of action for this cohort of KIs is the disruption of the cellular mitotic machinery, providing a key insight into both their therapeutic efficacy and potential toxicities.

## Discussion

Kinase inhibitors (KIs) have transformed cancer treatment, but their clinical utility is frequently undermined by a significant risk of cardiotoxicity, a major challenge in the field of cardio-oncology ^30^. The mechanisms driving these adverse events are complex, involving both on-target and off-target effects that disrupt essential pathways for cardiomyocyte health ^31^. A critical barrier to developing effective cardioprotective strategies has been the difficulty in distinguishing correlational molecular changes from true causal drivers ^32^. In this study, we developed KICDB, a comprehensive and interactive database, to address this gap by systematically integrating large-scale transcriptomic meta-analysis with robust causal inference through Mendelian randomization (MR). To our knowledge, KICDB is the first resource to provide the scientific community with an intuitive platform to explore the causal underpinnings of KI-induced cardiotoxicity.

A central strength of our study is the transcriptomics meta-analysis, which consolidated data from seven independent studies to define robust gene expression signatures for 26 different KIs. While numerous studies have provided invaluable insights into the mechanisms of individual KIs—such as sunitinib’s induction of ferroptosis and autophagic degradation ^33^, lenvatinib’s activation of endoplasmic reticulum stress ^14^, and osimertinib’s role in mitochondrial dysfunction ^34^—these focused approaches make it difficult to ascertain class-wide versus drug-specific effects. Our meta-analysis overcomes this limitation by identifying shared transcriptional programs, revealing, for instance, that a primary, common mechanism for many KIs is the disruption of the cellular mitotic machinery ^35^. This provides a systems-level view of the convergent pathways implicated in KI cardiotoxicity.

The most innovative aspect of KICDB is its integration of two-sample MR to infer causality, thereby moving beyond the correlational limitations of transcriptomic and observational studies. By leveraging genetic variants as instrumental variables, we formally tested the causal effects of kinase-related genes on 46 distinct cardiovascular outcomes. This large-scale analysis identified 26 significant causal associations, pinpointing novel molecular mediators of KI-induced cardiotoxicity. For example, our finding that genetically predicted higher expression of RNF13 is causally linked to an increased risk of primary cardiomyopathies provides a strong, data-driven hypothesis for future mechanistic studies ^36^. Conversely, the identification of genes such as TEK and JAK2 as having protective effects against conditions like atrial fibrillation indicates potential therapeutic targets for cardioprotection ^8^. These findings provide a critical layer of evidence that was previously missing in the field, offering a genetically validated starting point for biomarker and drug discovery.

While our study has its strengths, certain limitations must be acknowledged. First, the transcriptomic data are derived from in vitro human induced pluripotent stem cell-derived cardiomyocytes (hiPSC-CMs), which, despite their utility, may not fully recapitulate the complex physiology of the adult human heart ^37^. Second, while MR is a powerful tool for causal inference, it relies on assumptions that cannot be fully proven, such as the absence of horizontal pleiotropy ^38^. We sought to mitigate this by employing a suite of sensitivity analyses to ensure the robustness of our findings. Finally, the results presented in KICDB are intended to be hypothesis-generating and require further experimental validation to confirm the precise biological mechanisms.

Despite these limitations, KICDB represents a significant advance for the cardio-oncology community. The database empowers researchers to move beyond single-drug studies and explore systems-level patterns of cardiotoxicity across a wide range of KIs. This integrated approach can accelerate the validation of predictive biomarkers and aid in the rational design of targeted cardioprotective strategies ^39^. As the arsenal of targeted cancer therapies continues to expand, resources like KICDB will be indispensable for ensuring that improvements in oncologic outcomes are not offset by an undue burden of cardiovascular harm.

In conclusion, KICDB constitutes a unique and valuable resource by integrating large-scale transcriptomic data with causal inference to systematically investigate the mechanisms of KI-induced cardiotoxicity. By making these complex datasets accessible and queryable through an intuitive web interface, our work provides a powerful platform to formulate novel mechanistic hypotheses, guide future research, and ultimately contribute to the development of safer cancer therapies.

## Ethics Statement

This research article uses previously published data. It does not report on new human patient data from a prospective or retrospective study conducted by the author for this manuscript; therefore, institutional review board approval was not specifically required for the preparation of this article.

## Acknowledgement

Artificial intelligence (AI) programs were utilized for code development, manuscript writing, and data reporting for this study.

## Author Contributions

Conceptualization: J.M. Wei, Y. Liu, J. Zhang.

Project administration: J.M. Wei

Methodology: M.Q. Wu, G.Y. Li

Data curation: X.Y. Zheng, H.F. Fu Software: J. Zhang.

Writing – Original Draft: J.M. Wei,

Writing – Review & Editing: J.Zhang, J.J.Lin

## Data availability statement

The complete, interactive database is freely and publicly accessible at https://zhang-lab-database.shinyapps.io/KICDB/. The R scripts used for data analysis and to build the Shiny application are available from the corresponding authors upon reasonable request.

## References

1. Jin Y, Xu Z, Yan H, He Q, Yang X, Luo P. A Comprehensive Review of Clinical Cardiotoxicity Incidence of FDA-Approved Small-Molecule Kinase Inhibitors. Front Pharmacol. 2020;11:891.

2. Ben Ami E, Demetri GD. A safety evaluation of imatinib mesylate in the treatment of gastrointestinal stromal tumor. Expert Opin Drug Saf. 2016;15:571–578.

3. Lamore SD, Kohnken RA, Peters MF, Kolaja KL. Cardiovascular Toxicity Induced by Kinase Inhibitors: Mechanisms and Preclinical Approaches. Chem Res Toxicol. 2020;33:125–136.

4. Ekram J, Rathore A, Avila C, Hussein R, Alomar M. Unveiling the Cardiotoxicity Conundrum: Navigating the Seas of Tyrosine Kinase Inhibitor Therapies. Cancer Control. 2024;31:10732748241285756.

5. Lee DH, Hawk F, Seok K, Gliksman M, Emole J, Rhea IB, Viganego F, Welter-Frost A, Armanious M, Shah B, Chavez JC, Pinilla-Ibarz J, Schabath MB, Fradley M. Association between ibrutinib treatment and hypertension. Heart. 2022;108:445–450.

6. Karaağaç M, Eryılmaz MK. Pazopanib-induced fatal heart failure in a patient with unresectable soft tissue sarcoma and review of literature. J Oncol Pharm Pract. 2020;26:768–774.

7. Piper-Vallillo AJ, Costa DB, Sabe MA, Asnani A. Heart Failure Associated With the Epidermal Growth Factor Receptor Inhibitor Osimertinib. JACC CardioOncology. 2020;2:119–122.

8. Santos LG, Roque R, Antunes Santos R, Neves C, Bonito N. Severe Reversible Heart Failure Enhanced by Sorafenib Treatment for Hepatocellular Carcinoma. Cureus. 2025;17:e76993.

9. Scott SS, Greenlee AN, Matzko A, Stein M, Naughton MT, Zaramo TZ, Schwendeman EJ, Mohammad SJ, Diallo M, Revan R, Shimmin G, Tarun S, Ferrall J, Ho TH, Smith SA. Intracellular Signaling Pathways Mediating Tyrosine Kinase Inhibitor Cardiotoxicity. Heart Fail Clin. 2022;18:425–442.

10. Gawli CS, Patil BR, Nagpure NR, Patil CR, Kumar A, Patel HM. Prevalence of osimertinib-induced cardiotoxicity in non-small cell lung cancer patients: a systematic review and meta-analysis. Lung Cancer. 2025;205:108629.

11. Chen Z, Ai D. Cardiotoxicity associated with targeted cancer therapies. Mol Clin Oncol. 2016;4:675–681.

12. Yang H, Qiu S, Yao T, Liu G, Liu J, Guo L, Shi C, Xu Y, Ma J. Transcriptomics coupled with proteomics reveals osimertinib-induced myocardial mitochondrial dysfunction. Toxicol Lett. 2024;397:23–33.

13. Gunaydin Akyildiz A, Boran T, Jannuzzi AT, Alpertunga B. Mitochondrial dynamics imbalance and mitochondrial dysfunction contribute to the molecular cardiotoxic effects of lenvatinib. Toxicol Appl Pharmacol. 2021;423:115577.

14. Wang S, Ji F, Gao X, Li Z, Lv S, Zhang J, Luo J, Li D, Yan J, Zhang H, Fang K, Wu L, Li M. Tyrosine Kinase Inhibitor Lenvatinib Causes Cardiotoxicity by Inducing Endoplasmic Reticulum Stress and Apoptosis through Activating ATF6, IRE1α and PERK Signaling Pathways. Recent Pat Anticancer Drug Discov. 2024;20:168–184.

15. Wang H, Wang Y, Li J, He Z, Boswell SA, Chung M, You F, Han S. Three tyrosine kinase inhibitors cause cardiotoxicity by inducing endoplasmic reticulum stress and inflammation in cardiomyocytes. BMC Med. 2023;21:147.

16. Yang Z, Wan J, Zhang X, Mei J, Hao H, Liu S, Yi Y, Jiang M, He Y. Baicalin reduces sunitinib-induced cardiotoxicity in renal carcinoma PDX model by inhibiting myocardial injury, apoptosis and fibrosis. Front Pharmacol. 2025;16:1563194.

17. Grabowska ME, Chun B, Moya R, Saucerman JJ. Computational model of cardiomyocyte apoptosis identifies mechanisms of tyrosine kinase inhibitor-induced cardiotoxicity. J Mol Cell Cardiol. 2021;155:66–77.

18. Edgar R, Domrachev M, Lash AE. Gene Expression Omnibus: NCBI gene expression and hybridization array data repository. Nucleic Acids Res. 2002;30:207–210.

19. Love MI, Huber W, Anders S. Moderated estimation of fold change and dispersion for RNA-seq data with DESeq2. Genome Biol. 2014;15:550.

20. Viechtbauer W. Conducting meta-analyses in R with the metafor. J Stat Softw. 2010;36:1–48.

21. Hemani G, Zheng J, Elsworth B, Wade KH, Haberland V, Baird D, Laurin C, Burgess S, Bowden J, Langdon R, Tan VY, Yarmolinsky J, Shihab HA, Timpson NJ, Evans DM, Relton C, Martin RM, Davey Smith G, Gaunt TR, Haycock PC. The MR-base platform supports systematic causal inference across the human phenome. Elife. 2018;7:e34408.

22. Elsworth B, Lyon M, Alexander T, Liu Y, Matthews P, Hallett J, Bates P, Palmer T, Haberland V, Davey G, 1 S, Zheng J, Haycock P, Gaunt TR, Hemani G. The MRC IEU OpenGWAS data infrastructure. bioRxiv. 2020;:2020.08.10.244293.

23. Watanabe K, Stringer S, Frei O, Umićević Mirkov M, de Leeuw C, Polderman TJC, van der Sluis S, Andreassen OA, Neale BM, Posthuma D. A global overview of pleiotropy and genetic architecture in complex traits. Nat Genet. 2019;51:1339–1348.

24. Verbanck M, Chen CY, Neale B, Do R. Detection of widespread horizontal pleiotropy in causal relationships inferred from Mendelian randomization between complex traits and diseases. Nat Genet. 2018;50:693–698.

25. Korotkevich G, Sukhov V, Sergushichev A. fgsea: Fast Gene Set Enrichment Analysis. bioRxiv. 2023;:1–29.

26. Chang W CJAJSCSBXYAJMJDABB. shiny: Web Application Framework for R. R Packag. . 2025. Available at https://github.com/rstudio/shiny.

27. Chang W. shinydashboard: Create Dashboards with “Shiny”. https://CRAN.R-project.org/package=shinydashboard. R Packag. version 0.7. . 2015. Available at https://cran.r-project.org/package=shinydashboard.

28. Sievert C. Interactive Web-Based Data Visualization with R, plotly, and shiny. Chapman and Hall/CRC; 2020. doi:10.1201/9780429447273.

29. Almende B V., Thieurmel B. visNetwork: Network Visualization using “vis.js” Library. Cran. . 2016. Available at https://cran.r-project.org/web/packages/visNetwork/index.html.

30. Blaes A, Nohria A, Armenian S, Bergom C, Thavendiranathan P, Barac A, Sanchez-Petitto G, Desai S, Zullig LL, Morgans AK, Herrmann J. Cardiovascular Considerations After Cancer Therapy: Gaps in Evidence and JACC: CardioOncology Expert Panel Recommendations. JACC CardioOncology. 2025;7:1–19.

31. Gao M, Cheng Z, Yan W, Zhang Z, Zhang L, Geng H, Xu Y, Li C. Tyrosine kinase inhibitors induce cardiotoxicity by causing Ca(2+) overload through the inhibition of phosphoinositide 3-kinase activity. Biochem Biophys Res Commun. 2025;771:152027.

32. Wan Z, Wang C, Luo S, Zhu J, He H, Hao K. Bridging the Gap Between hiPSC-CMs Cardiotoxicity Assessment and Clinical LVEF Decline Risk: A Case Study of 21 Tyrosine Kinase Inhibitors. Pharmaceuticals (Basel). 2025;18. doi:10.3390/ph18040450.

33. Pan Z, Zhu L, Wang X, Huangfu N, Su P, Yang F, Fu X, Pu L, Fu Q, Chen J, Cui H, Liang P, Shen J. Excessive autophagic degradation of MYLK3 causes sunitinib-induced cardiotoxicity. Autophagy. 2025;:1– 20.

34. Kondo M, Nakamura Y, Kato Y, Nishimura A, Fukata M, Moriyama S, Ito T, Umezawa K, Urano Y, Akaike T, Akashi K, Kanda Y, Nishida M. Inorganic sulfides prevent osimertinib-induced mitochondrial dysfunction in human iPS cell-derived cardiomyocytes. J Pharmacol Sci. 2024;156:69–76.

35. Zhang H, To KKW. Serum creatine kinase elevation following tyrosine kinase inhibitor treatment in cancer patients: Symptoms, mechanism, and clinical management. Clin Transl Sci. 2024;17:e70053.

36. Bak M, Park H, Lee S-H, Lee N, Ahn M-J, Ahn JS, Jung HA, Park S, Cho J, Kim J, Park S-J, Chang S-A, Lee S-C, Park SW, Kim EK. The Risk and Reversibility of Osimertinib-Related Cardiotoxicity in a Real-World Population. J Thorac Oncol Off Publ Int Assoc Study Lung Cancer. 2025;20:167–176.

37. Gartrell J, Mulrooney DA, Hsu C-W, Pan H, Federico SM, Pappo AS, Helmig S, Meredith CJ, Pease P, Christensen AM, Beasley G, Bishop MW. Acute cardiotoxicity in pediatric and adolescent patients with solid tumors treated with tyrosine kinase inhibitors. Cancer. 2025;131:e35940.

38. Patel JN, Singh J, Ghosh N. Bruton’s tyrosine kinase inhibitor-related cardiotoxicity: The quest for predictive biomarkers and improved risk stratification. Oncotarget. 2024;15:355–359.

39. Deng Z, Xiao S, He Y-Y, Guo Y, Tang L-J. Sorafenib-induced cardiovascular toxicity: A cause for concern. Chem Biol Interact. 2025;410:111388.

